# The senescence regulator S40 family members from *Caragana intermedia* and *Arabidopsis thaliana* inhibit leaf senescence via promoting cytokinins synthesis

**DOI:** 10.1101/2022.04.08.487715

**Authors:** Tianrui Yang, Minna Zhang, Qi Yang, Kun Liu, Jiaming Cui, Jia Chen, Yufan Ren, Yunjie Shao, Ruigang Wang, Guojing Li

## Abstract

Leaf senescence is regulated by both endogenous hormones and environmental stimuli in a programmed and concerted way. The members of the S40 family have been reported to play roles in leaf senescence. Here we report that overexpression of an S40 family member from *Caragana intermedia, CiS40-11*, delayed the leaf senescence. Phylogenetic analysis revealed that the CiS40-11 protein had the highest identity with AtS40-5 and AtS40-6 of *A. thaliana. CiS40-11* was highly expressed in leaves and was down-regulated after dark treatment. The subcellular localization analysis showed that CiS40-11 was a cytoplasm-nucleus dual-localized protein. Leaf senescence was delayed in *CiS40-11* transgenic *A. thaliana* or by its transient expression in *C. intermedia*. Transcriptomic analysis and endogenous hormones assay revealed that CiS40-11 inhibited leaf senescence via promoting the biosynthesis of cytokinins, through blocking *AtMYB2* expression in *CiS40-11* overexpression lines. Furthermore, in the *ats40-5a* and *ats40-6a* mutants, *AtMYB2* expression was increased and their leaves exhibited a premature senescence phenotype. Our results show that CiS40-11 (and its orthologs, AtS40-5 and AtS40-6) promoted cytokinin synthesis by inhibiting the expression of *MYB2* and releasing its negative regulation on the expression of *IPTs* to inhabit leaf senescence.

**Highlight:** The senescence regulator *S40* family members *CiS40-11, AtS40-5* and *AtS40-6* are induced by light and inhibition leaf senescence by promoting cytokinin synthesis.

## Introduction

As the organ of photosynthesis, leaves are the major place where plants exchange materials and energy with the environment, and senescence is the last stage during plant development. When leaf senescence occurs, chloroplast grana, stroma, and plasma membrane are changed, diphosphoribulose carboxylase and chlorophyll a/b binding protein in chloroplast start to degrade, photosynthesis reduced, leaves turn yellow (Lim et al., 2007). Both exogenous environmental stimuli such as water, light, temperature and endogenous plant hormones and developmental stage can influence leaf senescence process, phytohormones as the core elements can integrate environmental and developmental factors from both hormone level and signal transduction regulation (Lim et al., 2007).

Inhibition of leaf senescence by cytokinin had been discovered for decades. After application of exogenous cytokinins, endogenous cytokinins content changed and the components of cytokinin signaling pathway altered, which all affected leaf senescence process (Gan et al., 1995; Kieber et al., 2014). In tobacco, the cytokinin synthesis gene, *IPT* (*Isopentenyl Transferase*), driven by the promoter of *SAG12* (*Senescence Associated Gene12*), significantly delayed leaf senescence (Gan et al., 1995). The transcription factor MYB2 suppressed the expression of *IPT1, IPT4, IPT5, IPT6*, and *IPT8* in *A. thaliana*. In the mutant *myb2-1*, the bioactive cytokinins significantly increased and the mutant exhibited a stay-green phenotype (Guo et al., 2011). In *Oryza sativa*, bioactive cytokinins increased by mutation of the *CKX* (Cytokinin Oxidase) gene, and the anti-senescence phenotype of the *osckx11* mutant was enhanced (Zhang et al., 2021). Cytokinin receptor AHK3 was a histidine kinase in *Arabidopsis*. In its loss-of-function mutant, the leaves showed premature senescence, and the senescence process of the double mutant *ahk2 ahk3* did not slow down even after applying exogenous cytokinin (Riefler et al., 2006). Mutation of *CRF6* (Cytokinin Response Factor 6) resulted in early senescence in the mutant *crf6*, indicating that CRF6 negatively regulated the leaf senescence process in a feedback way (Zwack et al., 2013).

The synthesis of cytokinins is an important way of regulating the content of active cytokinins in plants. The pathway of cytokinins synthesis was summarized as below: The isoprene group of DMAPP (Dimethylallyl Diphosphate) or DMAPP precursor was transferred to the adenine ring at the N6 position of ATP/ADP (Adenosine Triphosphate/Adenosine diphosphate) by IPT, and generating iPRTP/iPRDP (Isopentenyladenosine-5’-triphosphate/5’-diphosphate)(Kasahara et al., 2004), then iPRTP/iPRDP was transformed into cytokinin ribonucleic acid by CYP735A (Cytochrome Oxidase Protein735A), and ultimately was hydrolyzed to bioactive tZ (trans-Zeatin) by hydrolase LOG (Lonely Guy). And in the cytokinin synthesis pathway of *A. thaliana*, IPTs (1-9) were rate-limiting enzymes (Golovko et al., 2002; Kamada-Nobusada et al., 2009). In addition, the bioactive cytokinin content was also regulated by conversion into inactive forms through binding to sugars such as glucose, or via oxidative catabolism by CKXs (Werner et al., 2006).

Cytokinin signal transduction pathway consists of histidine kinase receptors AHKs (2-4), AHPs (Arabidopsis Histidine-containing Phosphor Transfer Proteins), and ARRs (Arabidopsis Response Regulators) (Ishida et al., 2008). At the terminal of leaf development stages, the expression level of transcription factor MYB2 increased and suppressed the expression of *IPTs*, while the expression level of *CKXs* increased, leading to the bioactive cytokinin level decreased. Next, the autophosphorylation of cytokinin receptor AHK3 was inhibited, and the phosphorylation signal transduction pathway of AHK3→AHPs→ARRs was blocked. Then the transcriptional activity of type B ARRs could not be activated, and the expression of cytokinin responsive genes, genes encoding extracellular transferases, and hexose transporter encoding genes that located downstream of type B ARRs were decreased as well, resulting in pool-source balance changed and *SAGs* expressed, and leaves eventually showed senescence phenotype (Lim et al., 2007).

The senescence regulator (SnR) family was also known as the S40 family, it had been named DUF584 previously (Fischer-Kilbienski et al., 2010). The number assigned to S40 family in the Pfam database is PF04520. It has a wide distribution in plants, and the family members have the conserved GRXLKGR(D/E)(L/M)XXXR(D/N/T)X(I/V)XXXXG(F/I) sequence at the carboxyl terminus (Fischer-Kilbienski et al., 2010). Becker et al. firstly found that the transcript level of *HvS40* increased significantly with leaf senescence in *Hordeum vulgate*, and thus initiated the study of S40 family in the regulation of leaf senescence process (Becker et al., 1993). Most members of the S40 family in *A. thaliana* were induced by dark treatment. There are 17 members of the S40 family in *O. sativa*, and the expression level of most genes was increased with natural senescence progression, except for *OsS40-4, OsS40-5, OsS40-7, OsS40-9, OsS40-14* and *OsS40-1* (Fischer-Kilbienski et al., 2010; Zheng et al., 2019). HvS40 localized in the nucleus, and it was abundantly expressed in mesophyll cells of the senescent leaves and was induced by SA and MeJA, the transcription factor WHIRLY1 inhibited the expression of *HvS40* (Krupinska et al., 2002; Krupinska et al., 2014). AtS40-3 belonged to S40 subfamily I and also localized in the nucleus, in its constitutively active mutant *s40-3a*, the delayed senescence phenotype was significantly enhanced (Fischer-Kilbienski et al., 2010). Furthermore, the S40 family was associated with seed germination, and the expression of *AtS40*.*4* was found to be induced by ABA. Knockout mutants of *AtS40*.*4* were hypersensitive to ABA treatment and on the contrary, its overexpression lines were insensitive to ABA at the seeds germination stage (Shi et al., 2021).

*Caragana intermedia* belongs to the *Caragana Fabr*., Leguminosae, and is a perennial shrub that has a strong ability to tolerate abiotic stresses (Xiao et al., 2003), but there is little research on leaf senescence regulation of *C. intermedia*. In addition, although the S40 family plays important roles during the leaf senescence process, the regulatory mechanism of this family remains unclear. In this study, *CiS40-11* was obtained from *C. intermedia*, and leaf senescence was delayed by both stable overexpression of *CiS40-11* in *A. thaliana* and by transient expression in *C. intermedia* seedlings. Combining transcriptome sequencing and endogenous phytohormones content detection, we found that the expression level in cytokinins synthesis genes increased significantly in *CiS40-11* overexpression lines, which led to an increase in endogenous cytokinins content in leaves and an enhancement of anti-senescence ability ultimately. While the two mutants of the *CiS40-11* orthologs in *A. thaliana* showed the contrary phenotype. Our work provides a new molecular mechanism of S40 family in leaf senescence regulation of plants.

## Materials and Methods

### Plant growth conditions and treatments

The wild-type (Columbia-0) and the transgenic lines were grown on 1/2 MS medium or a 1:3 mixture of peat soil and vermiculite under long-day conditions (22 °C, 16-h-light/8-h-dark). For the cytokinin treatment, seeds of wild type *A. thaliana* were sterilized and sown on 1/2 MS containing 1% agar powder, and supplemented with 2.5μM of 6-BA or with same volume of ddH_2_O (used as control). Seeds of the *CiS40-11* overexpressed *A. thaliana* were also sterilized and sown on 1/2 MS containing 1% agar powder. When the seedlings had grown for one week, the wild type that was either treated with 6-BA or under normal growth condition, as well as the *CiS40-11* overexpression lines were transferred to the same Petri dish and photographed.

Seeds of *C. intermedia* were collected from Wulanchabu City, Inner Mongolia Autonomous Region, China (41.44N, 111.69E), and sown in the 1:3 mixture of peat soil and vermiculite in a greenhouse (22 °C, 16-h-light/8-h-dark). Four weeks seedlings were subjected to dark treatment: Briefly, seedlings were placed in a 22 [incubator (RUMED) without light to grow for the desired time, and were observed every 2 days and samples were taken accordingly. Each sample contains three seedlings and was mixed evenly.

### Phylogenetic analysis and multiple sequence alignment

The *CiS40-11* sequence was selected from the drought-treated RNA-seq database of *C. intermedia* (SRA accession number: SRP121096) (Wan et al., 2018). The protein sequences of S40 members in *A. thaliana* were downloaded from the TAIR database (http://www.arabidopsis.org/). The phylogenetic analysis was conducted using MEGA7.0 with the neighbor-joining (NJ) method and 1000 bootstrap replications. Protein sequences alignments were carried out via the DNAMAN software.

### Construction and identification of plant materials

Full-length CDSs of *CiS40-11* was cloned into the expression vector pCanG-HA and pCAMBIA1302 using In-Fusion® HD Cloning Kit (TaKaRa, Japan). All generated binary plasmids were confirmed by enzyme digestion and sequencing validation, the recombinant vectors *35S*::*HA*-*CiS40-11* was used for phenotypic experiments, and the recombinant vectors *35S*::*CiS40-11-GFP* was used for subcellular localization observation. Ultimately, the binary plasmids were transformed into *Agrobacterium tumefaciens* strain GV3101 by electroporation used for stable transformation in *A. thaliana* and transient transformation in *C. intermedia* later. The mutants *ats40-5a* (Stock Number: GABI_331G02) and *ats40-6a* (Stock Number: SALK_110898) were obtained from ABRC (Arabidopsis Biological Resource Center) and were confirmed by PCR and qRT-PCR. Gene cloning and the mutants identification primers used were listed in Supplementary Tab. S3.

### Dark treatment of transiently transformed *C. intermedia*

*Agrobacterium*-mediated transient gene expression in *C. intermedia* mainly followed the procedure established by Liu et al. (Liu et al., 2019). In brief, plants were grown in the soil for about 3 weeks with sufficient watering. When the OD_600_ value of GV3101 reached 1.3-1.5, centrifuged the cell culture at 5000-6000 rpm for 10 minutes at 4□. Then re-suspended and diluted the cell pellet until the OD600 was 0.7-0.8 with 1/2MS medium (containing 100 μmol/L acetosyringone). Then resting the cell suspension at room temperature for 3-4h and mixing every half hour. Added 0.001% Silwet L-77 in the cell suspension before injection. Finally, used 1mL disposable syringes without needles to inject the evenly mixed cell suspension into the abaxial side of leaves. Two days after infiltration, the seedlings were subjected to dark treatment, and the seedlings infiltrated with pCanG-HA empty vector were used as control. When the seedlings showed senescence phenotype, photographs were taken immediately, and the yellow/green leaf ratio was calculated and the total chlorophyll content was measured. All the experiments were carried out with three biological replicates.

### RNA extraction and real time RT-PCR assay

Quantitative real-time polymerase chain reaction (qRT-PCR) was performed according to Yang et al. (Yang et al., 2018) The total RNA extraction kit (Tiangen, China) was used to extract total RNA. The RNA quantity and concentration were evaluated by agarose gel electrophoresis and Quawell micro volume spectrophotometer (Q5000, USA). Next, 1 μg of total RNA was reverse transcribed into cDNA using PrimeScript™ II 1st Strand cDNA Synthesis Kit (TaKaRa, Japan). qRT-PCR was performed with Roche LightCycler 480 Real Time PCR system (Roche, Switzerland) using LightCycler 480 SYBR Green Master (Roche, Switzerland) reagents. The cycling program was 95 °C for 30 s, followed by 40 cycles of 95 °C for 5 s, 60 °C for 30 s and 72 °C for 15 s. The melting curves were analyzed at 60-95 °C after 40 cycles. All qRT-PCR reactions were carried out with three technical replicates. The relative RNA transcript level of genes was calculated according to the 2^-ΔCT^ and 2^-ΔΔCT^ methods (Livak et al., 2001). *AtEF1*□ or *CiEF1*□ were chosen as the reference genes for the qRT-PCR analysis according to the plant materials (Yang et al., 2014). Primers used for qRT-PCR are listed in Supplementary Tab. S3. Each treatment was repeated three times independently.

### Subcellular localization

The T2 seeds of *35S*::*CiS40-11-GFP* transgenic lines grown on 1/2 MS medium and were selected with 13 mg/L hygromycin B, and the root tips of 10-day old positive seedlings were observed under the laser scanning confocal microscope (Zeiss LSM 510) with 488 nm wavelength.

### Transcriptome sequencing

Five-week-old wild type and the OE35 line were selected for transcriptome sequencing, with three replicates for each sample. RNA-seq was carried out by Majorbio Company (Shanghai, China). High-quality RNA samples (OD260/280=1.8∼2.2, OD260/230≥2.0, RIN≥6.5, 28S:18S≥1.0, >1μg) were used to construct the sequencing library. Transcriptome library was constructed following TruSeqTM RNA sample preparation Kit from Illumina (San Diego, CA) using 1μg of total RNA and sequenced with the Illumina HiSeq xten/NovaSeq 6000 sequencer. Raw paired end reads were trimmed and quality controlled by SeqPrep (https://github.com/jstjohn/SeqPrep) and Sickle (https://github.com/najoshi/sickle). Clean reads aligned to reference genome using HISAT2 (http://ccb.jhu.edu/software/hisat2/index.shtml) (Kim et al., 2015). Mapped reads were assembled by StringTie (https://ccb.jhu.edu/software/stringtie/index.shtml?t=example) (Pertea et al., 2015). Transcript level was calculated according to the transcripts per million reads (TPM) method. KEGG pathway analysis was carried out by KOBAS (http://kobas.cbi.pku.edu.cn/home.do) (Shen et al., 2014).

### Cytokinins detection

Four-week-old wild type *A. thaliana* and the OE35 line were selected for cytokinins detection, each sample with three replicates. The assay was carried out by Applied Protein Technology Co. Ltd (Shanghai, China). The standard working solutions were made by standards diluted with aqueous methanol to a series of concentrations, and the standard curves were established by the isotope internal standard method. Grind the samples in liquid nitrogen, add 30μL standard working solutions and 1mL acetonitrile aqueous solution (1% FA) solution to 100±5mg sample, shock blending, extract at 4 [for 12h, centrifuge at 14000g for 20min, 800 μL supernatant was taken and was dried with nitrogen, dissolved again with 100μL acetonitrile aqueous solution (1:1, v/v), centrifuged at 14000g for 20min, then taken the supernatant into ultra high performance liquid chromatograph (Waters I-Class LC). Mobile phase: liquid A is 0.05% FA aqueous solution, liquid B is 0.05% FA acetonitrile, the column temperature was 45°C, the flow rate was 400 μL/min, and the injection volume was 2 μL. The liquid phase gradients were as follows: 0-10 min, liquid B was varies linearly from 2% to 98%; 10-10.1 min, liquid B was varies linearly from 98% to 2%; 11.1-13 min, liquid B maintained at 2%, and used QC samples for quality control and inspection. Mass spectrometry analysis was performed using 5500 QTRAP mass spectrometer (AB SCIEX), and ESI resource condition was as below: source temperature 500[, Ion Source Gas1(Gas1):45, Ion Source Gas2(Gas2):45, Curtain gas (CUR):30, ionSapary Voltage Floating (ISVF)-4500 V, used MRM mode detection of the ion pair. The peak area and retention time were extracted using Multiquant, the cytokinins content was calculated by standard curve.

## Results

### CiS40-11 belongs to the SnR family with four conserved motifs

Bioinformatics analysis showed that CiS40-11 was composed of 210 amino acids, with a molecular weight of 23.06kDa. There are 15 numbers of S40 family in *A. thaliana*. To compare the divergence of CiS40-11 with other S40 proteins during evolution, CiS40-11 and S40 family members of *A. thaliana* with full-length protein sequences were conducted a phylogenetic analysis by MEGA7.0. As shown in Fig.1A, CiS40-11 was clustered to the same subfamily with AtS40-5, AtS40-6 and AtS40-11. The most obvious feature of the S40 family is that its C-terminal has a highly conserved amino acid sequence GRXLKGR(D/E)(L/M)XXXR(D/N/T)X(I/V)XXXXG(F/I) (Fischer-Kilbienski et al., 2010), we named it as motif 4 (Fig.1B). When combined with the sequence logos of Pfam database (Fig.S1) and the multiple alignment of CiS40-11, AtS40-5, AtS40-6 and AtS40-11 protein sequences, it’s intriguing to find the other three conserved motifs, and they were designated as motif1 (E/D)(L/F)XEX(D/E)(V/I/L), motif2 SXPXX(V/I)PXW and motif3 (PPHXXXX(R/K) respectively (Fig.1B). It suggested that these four motifs might play essential roles when S40 numbers performed functions.

**Fig. 1.**
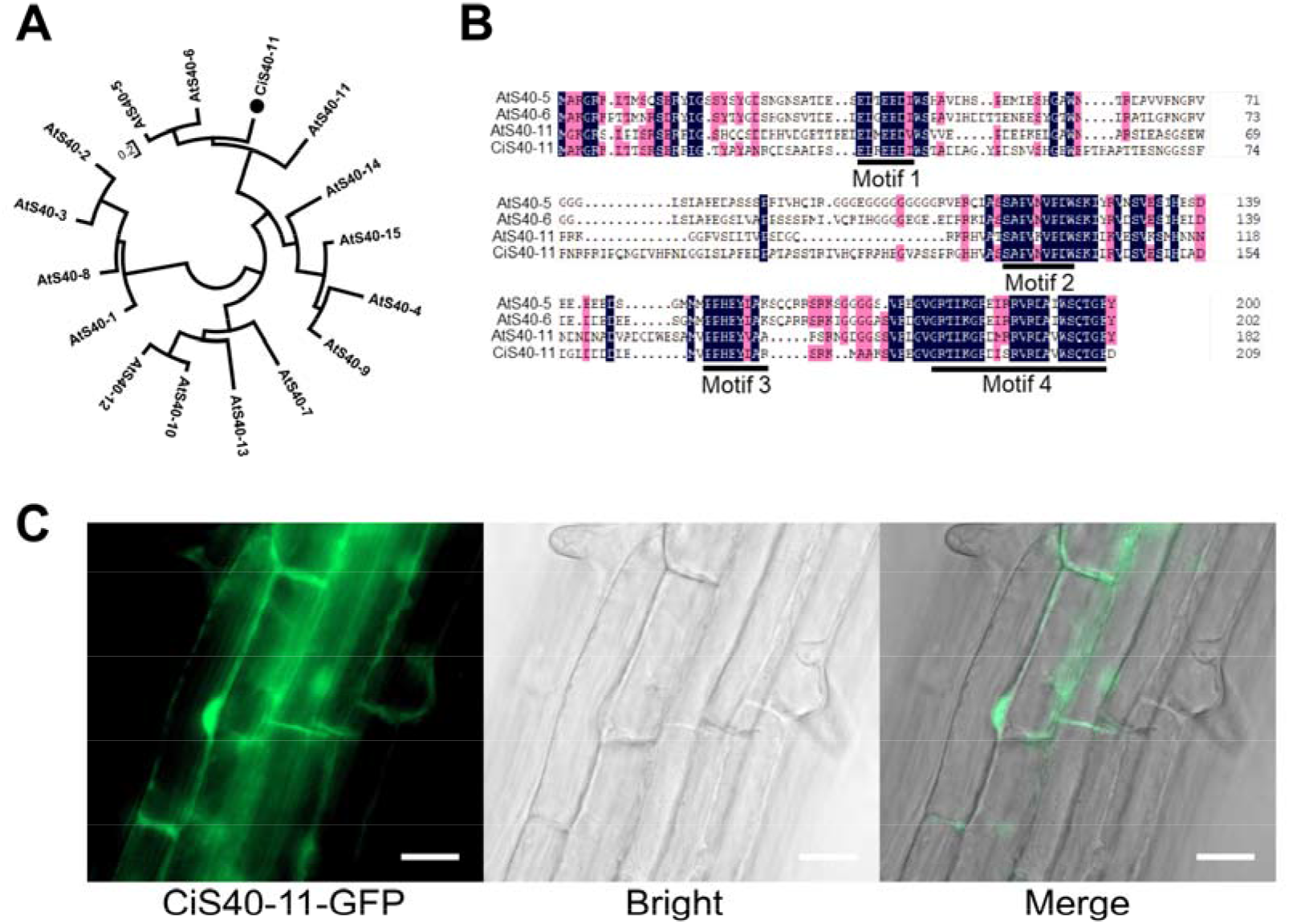
Phylogenetic analysis, protein sequences alignment and subcellular localization of CiS40-11. (A) Phylogenetic relationship of CiS40-11 and the S40 members in *A. thaliana*. The phylogenetic tree was generated with MEGA7.0 software by the neighbor-joining (NJ) method with 1,000 bootstrap replicates. CiS40-11 was marked with a black dot. (B) Amino acid alignment of CiS40-11 and its homologs of *A. thaliana*. Motifs (1-4) in amino acids sequence were indicated by underlines. Identical or similar amino acids were indicated with same color background. (C) Subcellular localization of CiS40-11. Root cells of *35S::CiS40-11-GFP* transgenic plants were used for observations of GFP signal by fluorescence microscope, bar=20μm.

### CiS40-11 is a cytoplasm-nucleus dual-localized protein

In order to confirm the subcellular localization of CiS40-11, we constructed fusion gene that the 3’ end of *CiS40-11* CDS (coding domain sequence) ligated with the *GFP* reporter gene and was driven by the CaMV (cauliflower mosaic virus) *35S* promoter. Based on the detected GFP signals in the root tip cells of transgenic *A. thaliana* seedling, we affirmed that CiS40-11 localized in both cytoplasm and nucleus (Fig.1C).

### The transcription of *CiS40-11* was down-regulated after dark treatment

To investigate the link between CiS40-11 and senescence, we detected the expression level of *CiS40-11* after dark treatment in *C. Intermedia*. Interestingly, unlike other reported S40 members, qRT-PCR results showed that *CiS40-11* was significantly down-regulated after dark treatment (Fig.2A). Moreover, we also detected the expression level of *CiS40-11* in roots, stems, leaves and the whole seedling, and found that *CiS40-11* was highly expressed in leaves (Fig.2B).

**Fig. 2.**
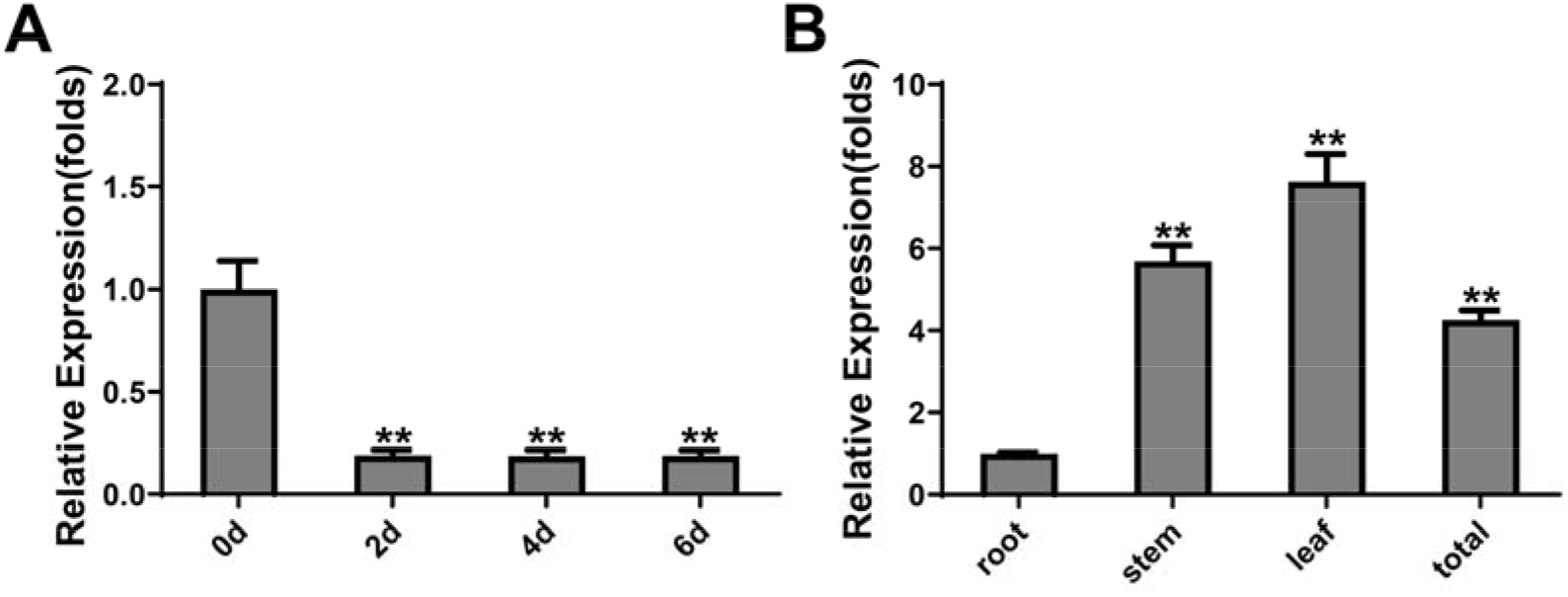
Gene expression patterns of *CiS40-11*. (A) Expression level analysis of *CiS40*-11 in 4 weeks old seedlings of *C. intermedia* with dark treatment. (B) Tissue specific expression analysis of *CiS40-11* in root, stem, leaves and the whole seedling of 4 weeks old *C. intermedia. CiEF1*α was used as the internal reference, and three independent biological replicates was carried out for each experiment. Three technical replicates were performed for each biological replicate during qRT-PCR analysis. ** represented P < 0.01.

### CiS40-11 delayed leaf senescence of both *C. intermedia* and *A. thaliana*

To uncover the function of *CiS40-11* during senescence process, we cloned the cDNA of *CiS40-11*from *C. intermedia* and constructed it with the pCanG-HA overexpression vector, driven by the CaMV *35S* promoter. Then the recombinant vector *35S::CiS40-11-HA* was transformed into *Agrobacterium tumefaciens* GV3101 for transient expression in *C. intermedia* seedlings and for stable transformation in *A. thaliana*. After 7 days of dark treatment, the leaves of control seedlings in *C. intermedia* that infiltrated with pCanG-HA empty vector showed a yellowish phenotype (Fig. 3A). Comparatively, the senescence degree of *CiS40-11* transient expression seedlings was slighter (Fig. 3A). The percentage of green and yellow-green leaves in *CiS40-11* transgenic seedlings was higher than that in the control (Fig. 3D). And the total chlorophyll content of *CiS40-11* transient expression seedlings (2.25 mg/g FW) was significantly higher than that of the control (2.02 mg/g FW) (Fig. 3D). Furthermore, we selected three transgenic *A. thaliana* lines with high *CiS40-11* expression levels for experimental analysis (Supplementary Fig. S1) and found that CiS40-11 also inhibited the senescence process in *A. thaliana*.

**Fig. 3.**
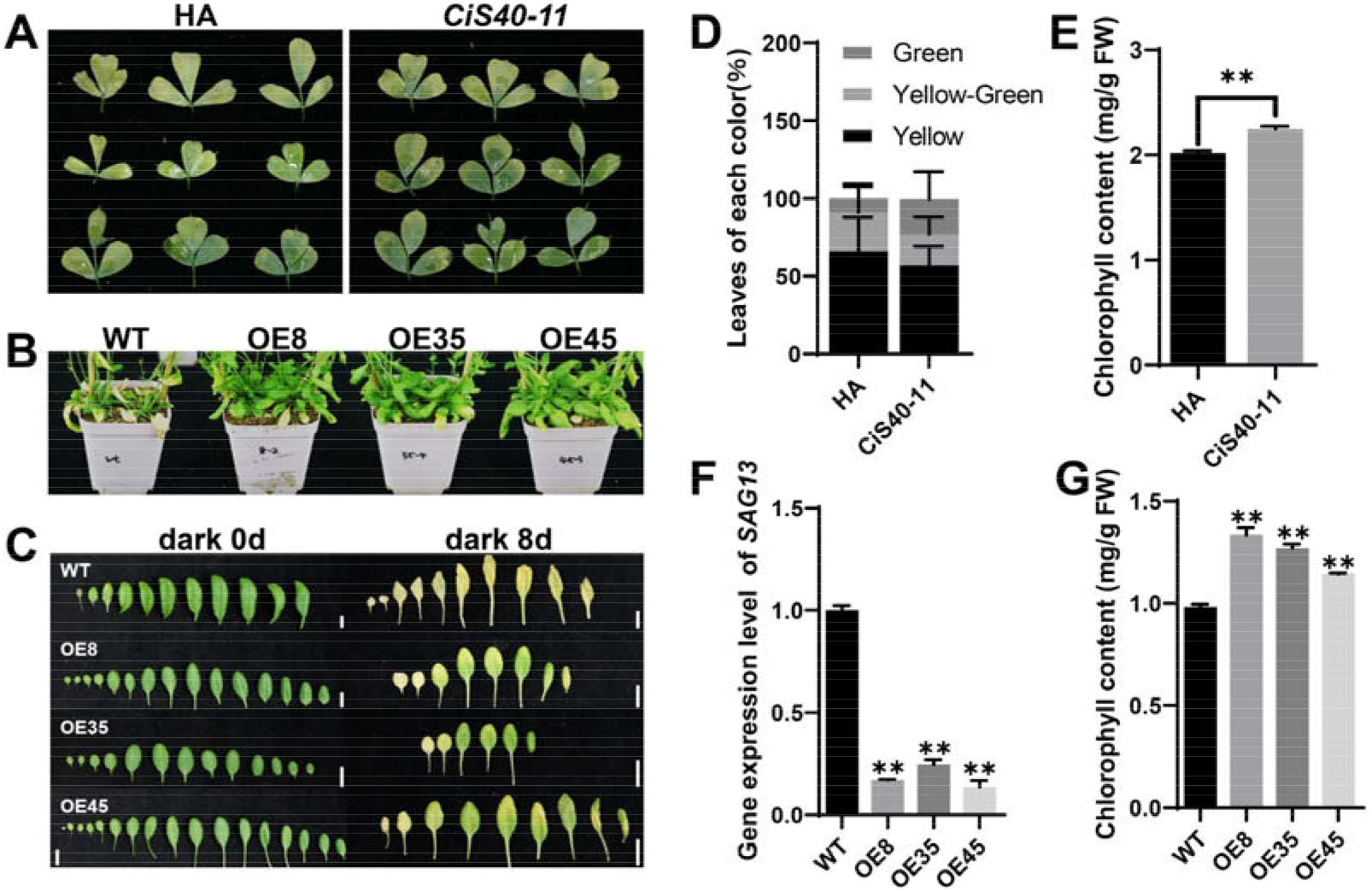
Senescence phenotype associated with *CiS40-11* transiently transformed *C. intermedia* seedlings and transgenic *A. thaliana*. (A) Dark-induced senescence phenotype of *CiS40-11* transiently transformed *C. intermedia* seedlings after 7 days darkness treatment. (B) Senescence phenotype of *CiS40-11* transgenic *A. thaliana* and wild type when the plant had grown about 8 weeks. (C) Dark-induced senescence phenotype of *CiS40-11* transgenic *A. thaliana* and wild type after darkness treatment for 8 days, bar = 1cm. (D) and (E) The statistical results of different color leaves percentage and the total chlorophyll content detection of *CiS40-11* transiently transformed *C. intermedia* seedlings after dark treatment. Each seedling was injected with three triple-emergent compound leaves, n=12. (F) *SAG13* expression level detection of 8-weeks old *CiS40-11* transgenic *A. thaliana* and wild type. The 5^th^, 6^th^, and 7^th^ rosette leaves were detected, n=3. (G) Total chlorophyll of the 5^th^, 6^th^, and 7^th^ rosette leaves of *CiS40-11* transgenic *A. thaliana* and wild type was detected after darkness treatment for 8 days, n=3. All the experiments were performed with three independent biological replicates; ** represented P < 0.01.

During the natural senescence process, the 8-weeks old *CiS40-11* overexpression lines were younger than the wild type (Fig. 3B), the expression level of senescence marker gene *SAG13* was declined significantly in *CiS40-11* overexpression lines compared with wild type (Fig. 3F). In the dark-induced senescence process, the senescence degree of *CiS40-11* overexpression lines was feebler than the wild type after 8 days of dark treatment. When most leaves of wild type turned yellow, the majority leaves of *CiS40-11* overexpression lines still stayed green (Fig. 3C), and the total chlorophyll content was higher in *CiS40-11* overexpression lines (Fig. 3G). These data indicated that CiS40-11 enhanced senescence resistance of both *C. intermedia* and *A. thaliana* leaves.

### The expression levels of cytokinin biosynthetic genes in *CiS40-11* overexpression lines significantly increased from transcriptome analysis

To further identify the effects of CiS40-11 on the leaf senescence process and to resolve the molecular mechanism, when *A. thaliana* had grown for five weeks and started showing senescence, we performed transcriptome sequencing with the *CiS40-11* overexpression line OE35 that had the highest expression level of *CiS40-11* in the transgenic *A. thaliana* (Supplementary Fig. S1A). The results showed that 1,125 genes were significantly up-regulated and 1,736 genes were significantly down-regulated (Supplementary Fig. S2).

The KEGG analysis of the *DEGs* (*Different Expression Genes*) showed that *DEGs* were significantly enriched in 11 pathways, and the enrichment gene numbers of plant hormone signal transduction were the largest and up to 57 genes (Fig. 4A). Therefore, we speculated that phytohormone levels might be changed in the overexpression lines. When functional annotation analysis of the top 200 *DEGs* genes with the most significant increase in the transcriptome was performed, we found that many of them were related to cytokinins (Supplementary Tab. S1). It had been well established that the leaf senescence was inhibited by cytokinin (Gan et al., 1995). To investigate whether the anti-senescence phenotype of *CiS40-11* overexpression lines was caused by endogenous cytokinins, we analyzed the expression levels of cytokinins biosynthetic genes in the transcriptome. The results showed that several *IPTs* expression was remarkably increased in the OE35 line, especially *IPT1* and *IPT3*, and the expression of *CYP35A2* was also increased (Fig. 4B). The qRT-PCR validation results were consistent with the transcriptome data (Fig. 4D). In addition, the expression levels of *CRF1-4, APRs*, and *LSU1-3*, which were known as the cytokinin responsive genes (Ohkama et al., 2002; Rashotte et al., 2006; Kim, 2016), were significantly increased in the OE35 line compared to the wild type (Fig. 4C). These results implied that the delayed leaf senescence of *CiS40-11* overexpression lines might be caused by high cytokinin content.

**Fig. 4.**
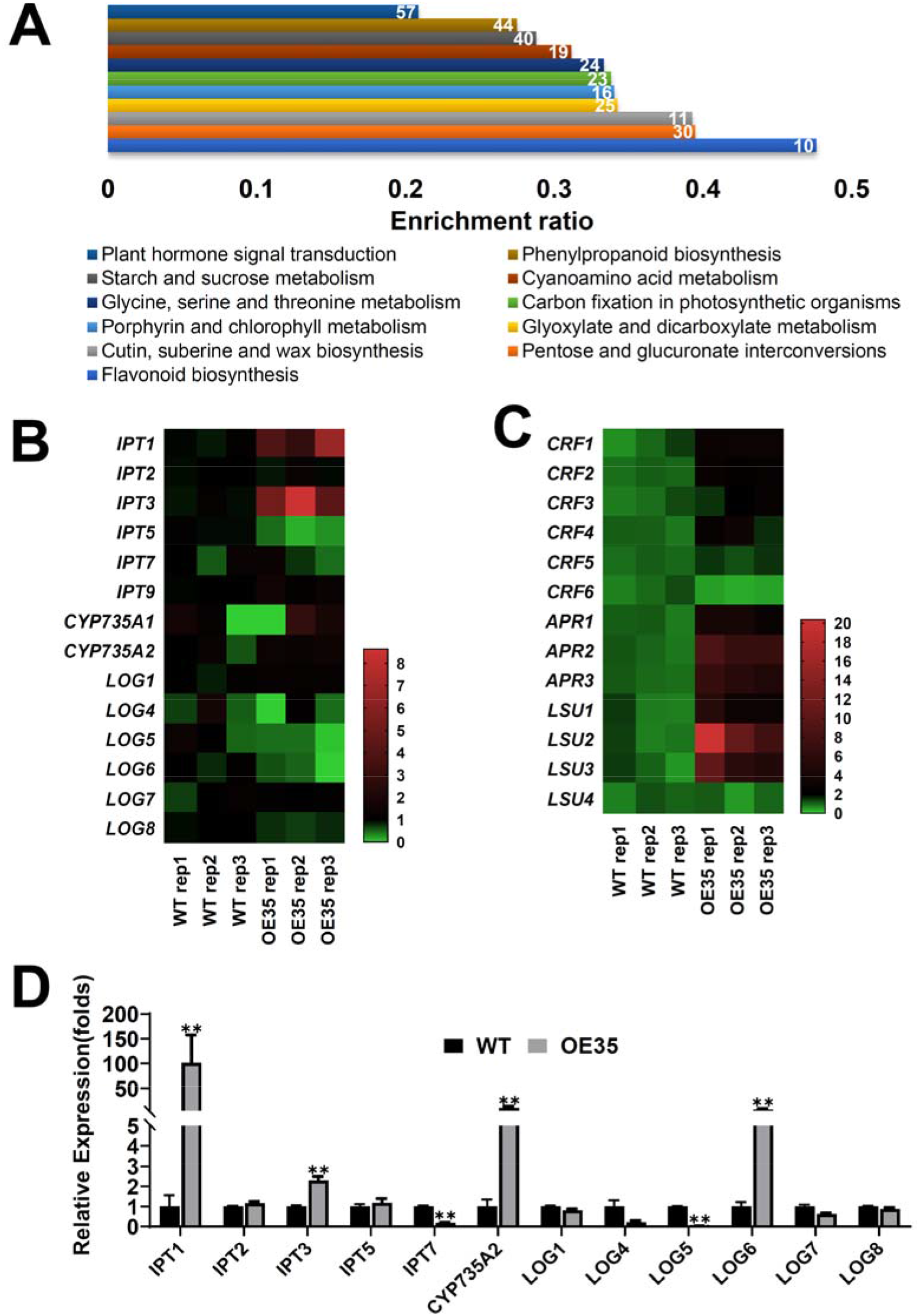
The transcriptome and cytokinin synthesis gene expression levels analysis of *CiS40-11* overexpression line in *A. thaliana*. (A) KEGG enrichment analysis of DEGs from the transcriptome. Metabolic pathways with P-value < 0.05 were shown, different color bars represent different metabolic pathways, the numbers within the bar represent the enrichment gene numbers. (B) and (C), the expression level of cytokinin synthesis genes and cytokinin responsive genes in transcriptome of the OE35 line and wild type, the color bar represents the relative expression values, red color represents the highest expression, and green color represents the lowest expression. (D) QRT-PCR validation of cytokinin synthesis genes in the OE35 line and wild type.

### Two active forms of cytokinins were increased in *CiS40-11* overexpression lines

Exogenously applied or increased endogenous cytokinins had been found to cause dwarfed, primary and lateral root development stunted, and leaf (Stenlid, 1982; Gan et al., 1996; Noodén et al., 1997; Ioio et al., 2012). The *CiS40-11* overexpression lines were dwarfed and the primary roots were shorter than that of the wild type, similar to the phenotype of the wild type after treating with 6-BA (a synthetic cytokinins analog) (Fig. 5A and Supplementary Fig. S3). And the total chlorophyll was remarkably increased in *CiS40-11* overexpression lines as well (Fig. 5B). We then measured the content of cytokinins, including iP, iPR, tZ, tZR, and czR in the OE35 line and wild type. The results showed that two active cytokinins forms, iPR, and tzR, increased obviously in the OE35 line (Fig. 5C). In summary, these data suggested that CiS40-11 inhibited the leaf senescence process by promoting *IPTs* expression and thus the synthesis increase of cytokinins in transgenic *A. thaliana*.

**Fig. 5.**
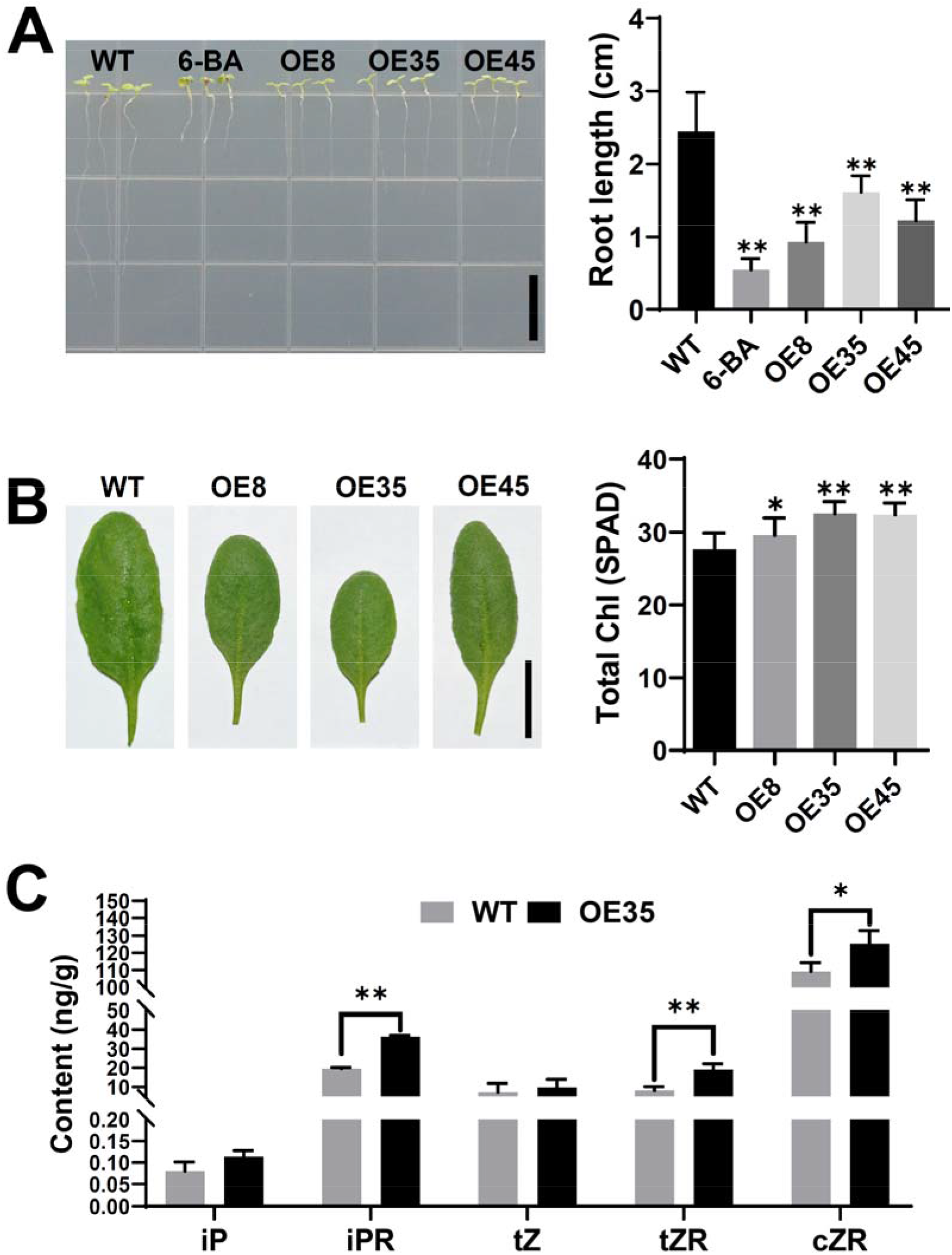
Cytokinins content detection and cytokinin-related phenotype of the *CiS40-11* overexpression lines. (A) Phenotype of the primary roots of seedlings of different genotypes or under 6-BA treatment and the average root length. Length of the primary roots was measured when the seedlings had grown for 7 days, n ≥ 36, ** P < 0.01, bar = 1cm. (B) The leaf morphology and the total chlorophyll content of leaves. The total chlorophyll of the 5^th^, 6^th^, and 7^th^ rosette leaves was detected by SPAD chlorophyll meter and the average value was calculated when plants of different genotypes had grown for 4 weeks, n ≥ 20, bar = 1cm. (C) Endogenous cytokinins content detection of OE35 line and wild type when plants had grown for 4 weeks, with three independent technical replicates. * P < 0.05; ** P < 0.01.

### Knockdown of the *CiS40-11* orthologs of *A. thaliana* accelerated leaf senescence

CiS40-11 had two close orthologs in *A. thaliana*, AtS40-5 and AtS40-6 (Fig. 1A). In order to exclude the effect of heterologous expression, the functions of AtS40-5 and AtS40-6 were investigated. Overexpression of either *AtS40-5* or *AtS40-6* in *A. thaliana* showed a similar phenotype as the *CiS40-11* transgenic lines (Supplementary Fig. S4), indicating they had similar functions. We then obtained *ats40-5a* and *ats40-6a*, the knockdown mutants of *AtS40-5* and *AtS40-6* respectively (Supplementary Fig. S5). And leaf senescence of both *ats40-5a* and *ats40-6a* mutants progressed significantly faster than that of wild type, when plants had grown about 6.5 weeks (Fig.6A). The total chlorophyll content was declined and the expression level of *SAG113* was increased in *ats40-5a* and *ats40-6a* as well (Fig.6A). In brief, mutants *ats40-5a* and *ats40-6a* showed the opposite phenotype compared with the *CiS40-11* overexpression line. We also examined the expression levels of *IPTs* in the mutants *ats40-5a* and *ats40-6a*, and found that the transcript of *IPT3* was reduced in both mutants (Fig. 6B).

**Fig. 6.**
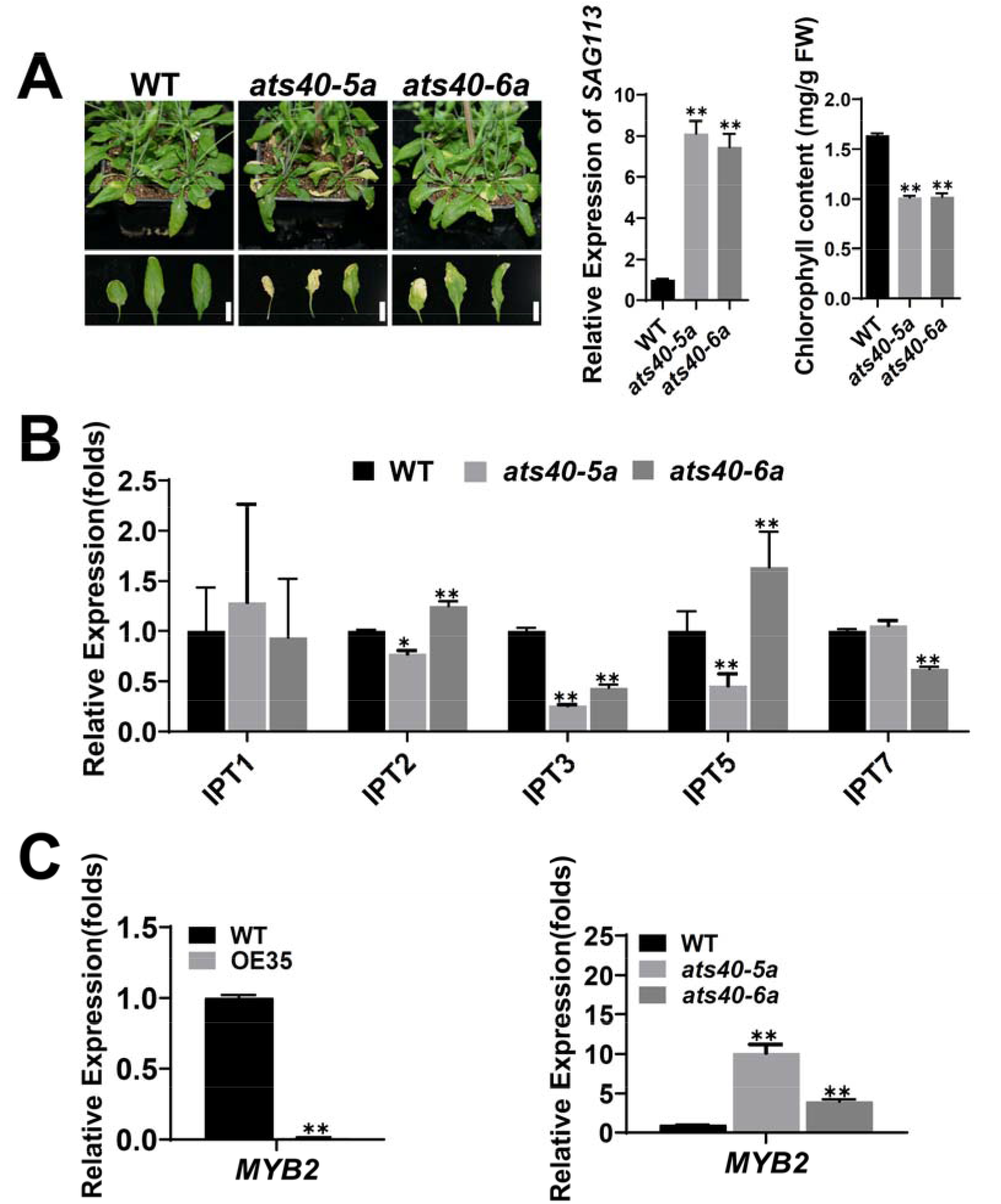
Senescence phenotype of *CiS40-11* ortholog mutants of *A. thaliana* and gene expression levels detection of *IPTs* and *AtMYB2* in mutants, the *OE35* line and wild type. (A) Senescence-related phenotype of the *ats40-5a* and *ats40-6a* mutant*s* and wild type, the histograms from left to right are results of *SAG113* expression level and total chlorophyll content detection in the *ats40-5a* and *ats40-6* mutants, bar = 1cm. (B) qRT-PCR validation of *IPTs* in the *ats40-5a* and *ats40-6* mutants. (C) qRT-PCR detection of *AtMYB2* expression in the OE35 line, the *ats40-5a* and *ats40-6a* mutants and wild type. When the plant had grown about 6.5 weeks and showed visible senescence phenotype, the 5^th^, 6^th^, and 7^th^ rosette leaves were selected and mixed to detect the relevant physiological indicators. All the experiments were carried out with three biological replicates, n ≥ 3; ** means P < 0.01.

### The expression of *AtMYB2* was enhanced in mutants *ats40-5a* and *ats40-6a*

*AtMYB2* abundantly expressed during leaf senescence and inhibited *IPTs* expression, leading to endogenous cytokinins content decrease, which accelerated the leaf senescence process eventually (Guo et al., 2011). To further determine whether CiS40-11 and its orthologs promoted *IPTs* transcription by suppressing the expression of *AtMYB2* during leaf senescence, we detected the expression level of *AtMYB2* in OE35 line, *ats40-5a* and *ats40-6a* mutants respectively. The expression of *AtMYB2* had dropped drastically in the OE35 line; on the contrary, its transcript level had elevated significantly in *ats40-5a* and *ats40-6a* mutants (Fig. 6C). Hence, we concluded that CiS40-11 and its orthologs promoted *IPTs* expression by suppressing *AtMYB2* at the transcriptional level.

## Discussion

Leaf senescence is intimate with phytohormones, no matter the initiation, progression, or the terminal phases of senescence, are all well concerted by plant hormones. The regulation of phytohormones on senescence could be considered as either positive or negative (Lim et al., 2007; Guo et al., 2021). Abscisic acid (ABA), ethylene, jasmonic acid/methyl jasmonate, and salicylic acid facilitated leaf senescence, while cytokinins, auxin, brassinosteroid, and gibberellins have been reported to postpone leaf senescence (Gan et al., 1995; Morris et al., 2000; Kim et al., 2011; Yu et al., 2011; Danisman et al., 2012; Zhang et al., 2012; Chang et al., 2013; Chen et al., 2017). In this study, we found CiS40-11 could inhibit the leaf senescence process by promoting the synthesis of the cytokinin in *A. thaliana* (Fig. 4D and 5B). And the cytokinins cross-talked with others phytohormones, for example, the endogenous cytokinins increase caused ABA levels to decrease (Jeon et al., 2010; Zhang et al., 2021). Woeste et al. demonstrated that cytokinins improved ethylene levels and suggested cytokinins increased ACS4 function by post-transcriptional regulation (Woeste et al., 1999). In the process of plant growth, cytokinin biosynthesis was reduced by auxin treatment, while treatment with cytokinins promoted auxin synthesis in young leaves (Takei et al., 2004; Jones et al., 2010). During the cell differentiation of root meristem growth phase, cytokinins and gibberellins showed antagonistic regulation of ARR1(Moubayidin et al., 2010). Thus, we speculated that during leaf senescence, CiS40-11 directly or indirectly influenced others hormones levels. However, how could CiS40-11 affect other phytohormones and the holistic hormonal regulatory network remained to be investigated in future.

*AtMYB2* expression was inhibited by CiS40-11 and its homologs, AtS40-5 and AtS40-6 (Fig. 6C). Though there were a few reports on the regulation of downstream genes expression by the S40 family in *A. thaliana*, such as AtS40-3 inhibited *WRKY53* expression (Fischer-Kilbienski et al., 2010), AtS40.4 regulated the transcription of *ABI* (*ABA Insentive*)*1-5* and *SnRK2*.*6* (Shi et al., 2021), but the regulatory mechanism of S40 family had remained elusive. There was a substantial proportion of S40 members localized in the nucleus. It had been shown that AtS40-3 and HvS40 did not contain a classical nuclear localization sequence, but they did have the PEST sequence which was typical for proteins with high turnover being degraded by the ubiquitin/proteasome system, and several proteins which had PEST were proved to be located in the nuclei (Fischer-Kilbienski et al., 2010; Lagrange et al., 2003). CiS40-11 was localized not only in the cytoplasm but also in the nucleus. Although there were no classic nuclear localization sequence were predicted by NLStradamus, we did find the PEST sequence in CiS40-11 (Supplementary Fig. S6). We speculated that during the leaf senescence stage, CiS40-11 might bind to the currently unknown transcription factor that regulated *MYB2* expression and then degraded it through the protease degradation pathway, this led to the transcription of *MYB2* decreased and promoted *IPTs* expression ultimately, so the synthesis of cytokinins process was enhanced and the leaf senescence process was suppressed.

In barley, WHIRLY1 is directly bound to the *HvS40* promoter and regulated *HvS40* expression (Krupinska et al., 2014). Interestingly, WHIRLY1 did not regulate *AtS40-3* expression but it suppressed *WRKY53* expression in *A. thaliana* (Miao et al., 2013). Jehanzeb et al. analyzed the promoter sequences of *S40* family numbers and discovered multiple binding sites or recognition sequences of WRKY, MYC, MYB, ABRE, Dof, and DRE/CRT transcription factors (Jehanzeb et al., 2017), and this indicated that *S40* family members might be regulated by these transcription factors in *A. thaliana*. Because the genome of *C. intermedia* has not been sequenced yet, we cloned the promoter of *CiS40-11* by genome-walking technology. Unfortunately, we did not obtain a suitable length of its promoter fragment. Thus, we analyzed the 2000bp promoter sequence upstream of the start codons of *AtS40-5* and *AtS40-6*, and found a large number of light-responsive elements in both promoters (Supplementary Tab. S2), we shaded the *Pro*_*AtS40-5*_::*GUS* and *Pro*_*AtS40-5*_::*GUS* transgenic *A. thaliana* leaves by wrapping them with tinfoil, and histochemical staining for β-glucuronidase (GUS) Jefferson(Jefferson et al., 1987) showed that the half leaf covered by the tinfoil was not stained at all, while the half leaf under normal light was stained obviously, indicating that light promoted the expression of *AtS40-5* and *AtS40-6* (Supplementary Fig. S7A). The *CiS40-11* expression level showed a significant decrease after dark treatment (Fig. 2B), we therefore inferred that light was an important factor in *CiS40-11* expression regulation. Then we restored lighting for the dark-treated *C. intermedia* seedlings and found that the expression level of *CiS40-11* increased gradually with the prolonged lighting time (Supplementary Fig. S7B). These results suggest that light is an important factor of promoting *CiS40-11, AtS40-5*, and *AtS40-6* expression. It was interesting to note that besides light-responsive elements, we also found many MYB recognition sequences in *AtS40-5* and *AtS40-6* promoters (Supplementary Tab. S2). Therefore, we could not exclude the possibility that MYB2 might also regulate *CiS40-11* and its homolog genes expression.

During leaf senescence process, it had been reported that the expression level of thousands of *SAGs* (*Senescence-associated Genes*) was up-regulated and caused the senescence syndrome (Guo et al., 2005; Woo et al., 2019; Li et al., 2020). To determine which downstream *SAGs* were directly regulated by the cytokinins in the leaf senescence pathway that CiS40-11 mediated, in combination with transcriptome analysis, we selected several *SAGs* with significantly decreased expression as candidate genes and validated with qRT-PCR (Supplementary Fig. S8). It showed that the expression levels of *SAG12, SAG102, SAG201* were decreased notably both in the transcriptome database and qRT-PCR validation with the *CiS40-11* overexpression line, and the expression level of *SAG12* decreased the most remarkably. Noticeably, *SAG12* was also significantly down-regulated in the *AtS40-3* overexpression mutant, *ats40-3a*, and WRKY53 promoted *SAG12* expression (Miao et al., 2004; Fischer-Kilbienski et al., 2010). Further analysis of the expression level of *WRKY53* in transcriptome showed that it was also down-regulated in the *CiS40-11* overexpression line. We speculated that AtS40-3 might also inhibit leaf senescence through the cytokinin-mediated pathway and WRKY53 might locate downstream of the cytokinin signaling pathway. In addition, although *SAG29* was significantly up-regulated in the transcriptomic data, the qRT-PCR test revealed that it was also significantly down-regulated actually. These *SAGs* would be the focus of our next research.

Hence, we proposed a working model on leaf senescence regulation: CiS40-11 (and its orthologs, AtS40-5 and AtS40-6) promoted cytokinin synthesis by inhibiting the expression of *MYB2* and releasing its negative regulation on the expression of *IPTs* in *A. thaliana* (Fig.7). Our evidence here provided the new research platform and genetic resources for further uncovering the action mechanism of the S40 family.

**Fig. 7.**
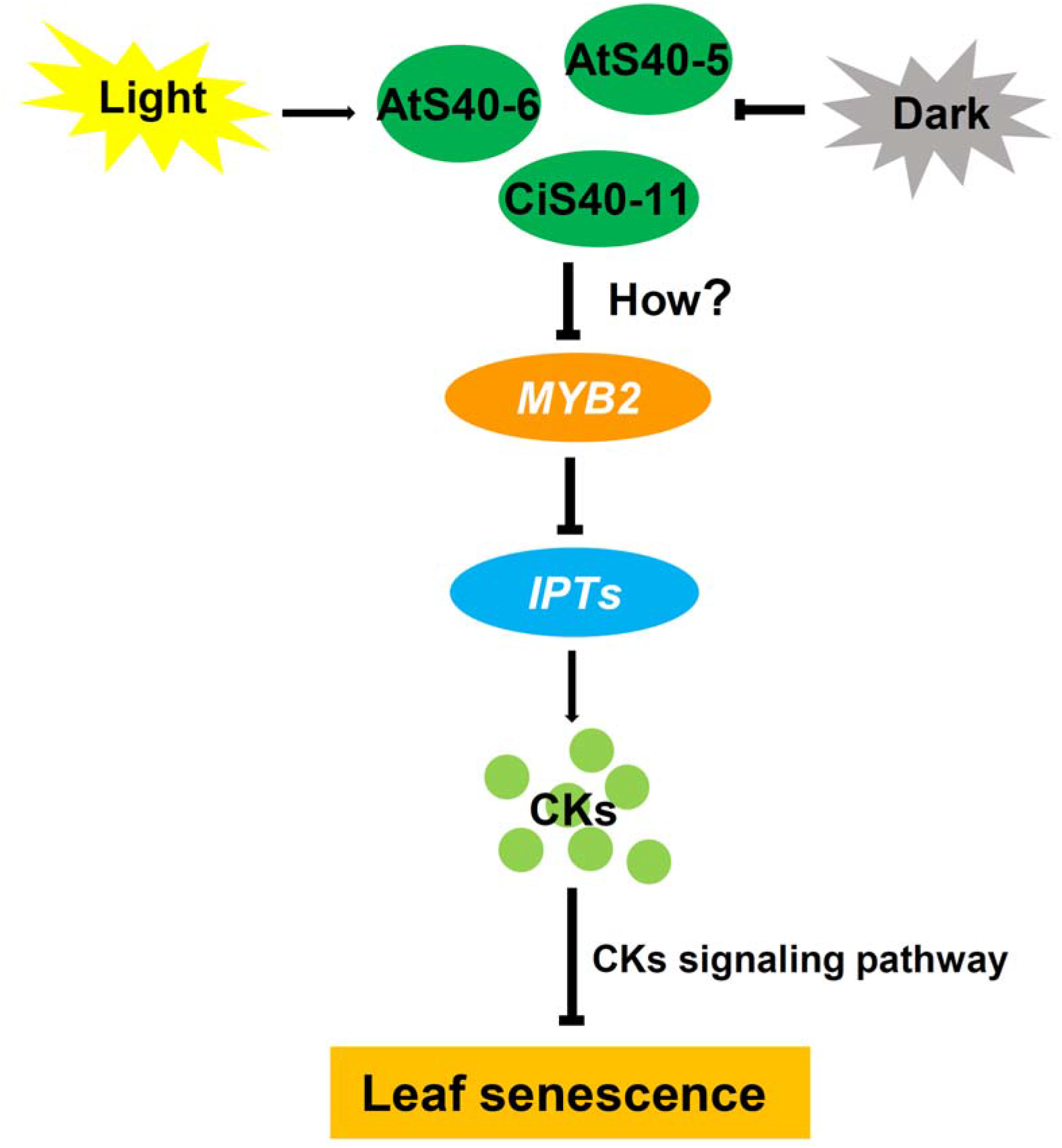
The regulation mechanism model of CiS40-11 and its orthologs on leaf senescence. CiS40-11 (and AtS40-5/6) increases endogenous cytokinins accumulation through inhibiting *AtMYB2* mediated signal transduction. Light promotes CiS40-11 (and AtS40-5/6) expression to inhibit *AtMYB2* expression, which in turn leads to an increased expression of *IPTs*, and results in the active form of cytokinins content increasing in plants, this ultimately retards leaf senescence process.

## Supplementary data

There are 8 Supplementary Figures and 3 Supplementary Tables in this study.

**Supplementary Figure S1**. The target gene expression level detection of the transgenic *A. thaliana*.

**Supplementary Figure S2**. The gene expression profiling from the transcriptome data of the OE45 line and wild-type.

**Supplementary Figure S3**. Phenotype of the primary roots of seedlings of different genotypes or under 6-BA treatment.

**Supplementary Figure S4**. The senescence phenotype of the *AtS40-5* and *AtS40-6* overexpression lines and their corresponding mutants *ats40-5a* and *ats40-6a*.

**Supplementary Figure S5**. The characterization of T-DNA insertion mutants *ats40-5a* and *ats40-6a*.

**Supplementary Figure S6**. The nuclear localization sequence and the PEST sequence prediction of CiS40-11.

**Supplementary Figure S7**. The histochemical staining of *Pro*_*AtS40-5*_::*GUS* and *Pro*_*AtS40-6*_::*GUS* transgenic *A. thaliana* and the *CiS40-11* expression level detection under light treatment.

**Supplementary Figure S8**. The expression level of some *SAGs* changed in the transcriptome and qRT-PCR validation in the OE35 line.

**Supplementary Table S1**. The top 200 genes significantly upregulated in the transcriptome of the OE35 line of CiS40-11 transgenic A. thaliana.

**Supplementary Table S2**. Analysis of cis-elements of AtS40-5 and AtS40-6 promoters in this study.

**Supplementary Table S3**. The primer sequences used in this study.

## Acknowledgments

We thank Dr. Qi Xie, Institute of Genetics and Developmental Biology, Chinese Academy of Science, kingly provided us the pCanG-HA vector.

## Author contributions

Guojing Li, Ruigang Wang and Tianrui Yang designed experiments. Tianrui Yang and Minna Zhang carried out experiments with the help of Kun Liu, Jiaming Cui, Jia Chen, Yufan Ren and Yunjie Shao. Tianrui Yang analyzed the transcriptome data with the help of Qi Yang. The manuscript was written by Tianrui Yang and Minna Zhang, Guojing Li and Ruigang Wang edited it. Tianrui Yang and Minna Zhang contributed equally to this work.

## Data Availability

The *CiS40-11* nucleotide sequence have been deposited at the GenBank database with the accession ON112076. The transcriptomic data have been deposited at the Sequence Read Archive (SRA) database with the accession SRP367210.

