## Supplement Figure S1 to S8 for "The senescence regulator S40 family members from *Caragana intermedia* and *Arabidopsis thaliana* inhibit leaf senescence via promoting cytokinins synthesis"

**Supplementary data:**

There are 8 Supplementary Figures and 3 Supplementary Tables in this study.

**Supplementary Figure S1.** The target gene expression level detection of the transgenic *A. thaliana*.

**Supplementary Figure S2.** The gene expression profiling from the transcriptome data of the OE45 line and wild-type.

**Supplementary Figure S3.** Phenotype of the primary roots of seedlings of different genotypes or under 6-BA treatment.

**Supplementary Figure S4.** The senescence phenotype of the *AtS40-5* and *AtS40-6* overexpression lines and their corresponding mutants *ats40-5a* and *ats40-6a*.

**Supplementary Figure S5.** The characterization of T-DNA insertion mutants *ats40-5a* and *ats40-6a*.

**Supplementary Figure S6.** The nuclear localization sequence and the PEST sequence prediction of CiS40-11.

**Supplementary Figure S7.** The histochemical staining of *Pro_AtS40-5_::GUS* and *Pro_AtS40-6_::GUS* transgenic *A. thaliana* and the *CiS40-11* expression level detection under light treatment.

**Supplementary Figure S8.** The expression level of some *SAGs* changed in the transcriptome and qRT-PCR validation in the OE35 line.

**Supplementary Table S1.** The top 200 genes significantly upregulated in the transcriptome of the OE35 line of CiS40-11 transgenic A. thaliana.

**Supplementary Table S2.** Analysis of cis-elements of AtS40-5 and AtS40-6 promoters in this study.

**Supplementary Table S3.** The primer sequences used in this study.

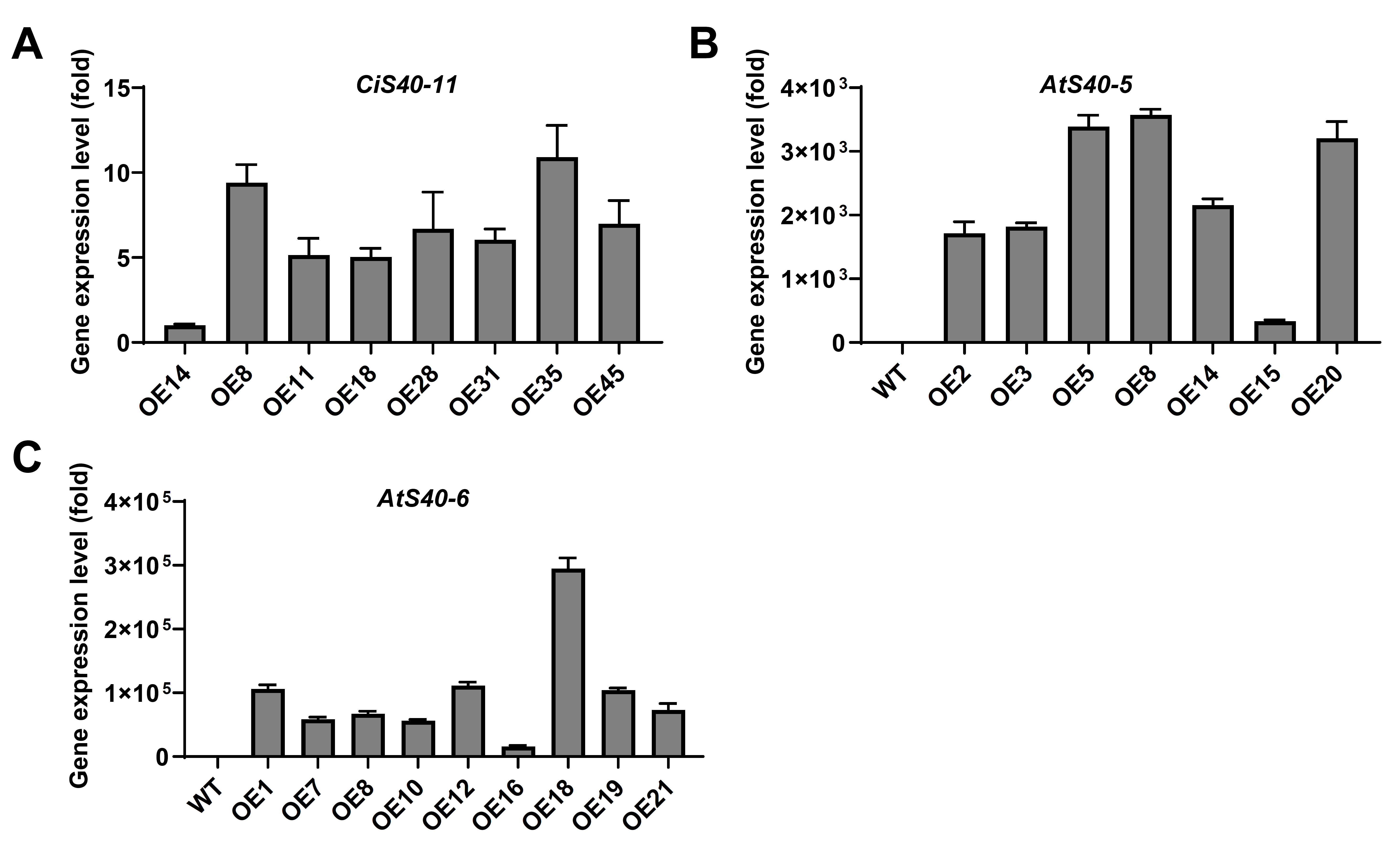

**Supplementary Figure S1.** **The target gene expression level detection of the transgenic *A. thaliana*.**

The transcript level of *CiS40-11* (A), *AtS40-5* (B) and *AtS40-6* (C) was detected by qRT-PCR in the correspondingly transgenic lines of *A. thaliana*. *AtEF1α* was taken as the internal reference gene and the expression value was calculated using 2^-ΔΔCT^ method. The OE14 was the positive control in (A), WT was the positive control in (B) and (C). Two independent biological replicates were performed, and three technical replicates for each sample were carried out.

**Supplementary Figure S2. The gene** **expression profiling from the transcriptome data of the OE45 line and wild-type.**

The heat map and the pie chart of gene expression profiling from the transcriptome analysis were shown here. The red, violet and white colors represented the differential expression genes (DEGs) that significantly up-regulated, down-regulated or unchanged respectively. The number 1, 2, and 3 indicated the replicate samples.

**Supplementary Figure S3. Phenotype of the primary roots of seedlings of different genotypes or under 6-BA treatment.**

Bar = 1cm, the treatment methods and plant growth conditions are shown in the legend of (Fig. 5A) and the Methods section.

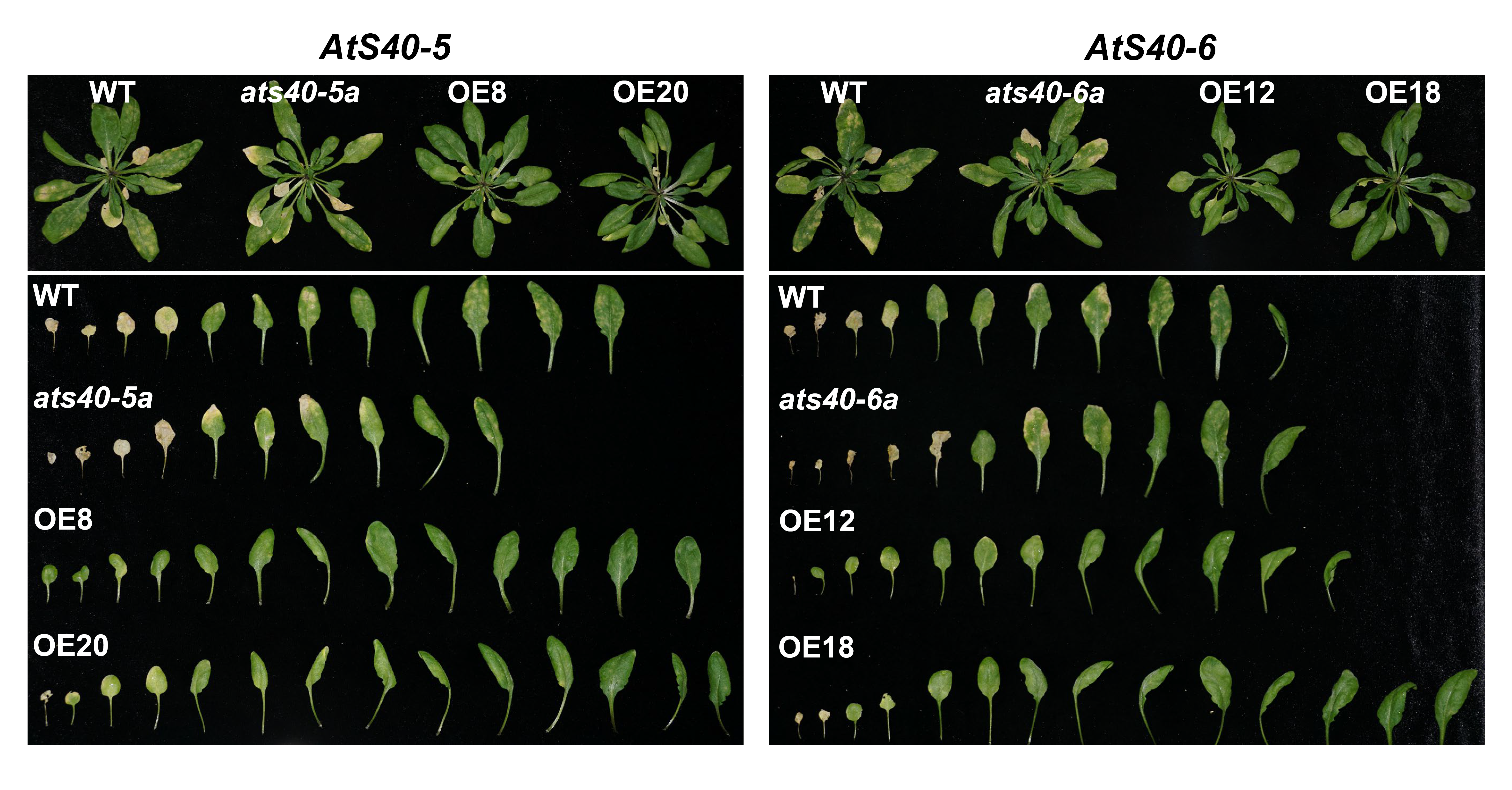

**Supplementary Figure S4.** **The senescence phenotype of the *AtS40-5* and *AtS40-6* overexpression lines and their corresponding mutants *ats40-5a* and *ats40-6a*.**

Plants were grown for about 6.5 weeks and then the photographs were taken with camera (Sony).

**Supplementary Figure S5.** **The characterization of T-DNA insertion mutants *ats40-5a* and *ats40-6a*.**

(A) was the schematic map of the T-DNA insertion sites in *ats40-5a* and *ats40-6a* mutants. The gray lines represented the UTR structure, the black arrows represented the exons, and the black triangles represented the T-DNA insertion sites. (B) was the electropherograms of the PCR identification of the homozygote mutants. The PCR products amplified by the left + right primers of *AtS40-5* (Lane 1), 8474 + the right primer of *AtS40-5* (Lane 2), the left + right primers of *AtS40-6* (Lane 3) and LBb1.3 + the right primer of *AtS40-6* (Lane 4) was shown respectively. 8474 and LBb3 was the left border primer of the T-DNA from both GABI and SALK vectors respectively. (C) was the transcript level of *AtS40-5* and *AtS40-6* in the mutants *ats40-5a* and *ats40-6a* with qRT-PCR detection respectively. *AtEF1α* as used as the internal reference gene and the expression value was calculated by 2^-ΔΔCT^ method, WT was the positive control. Two independent biological replicates were performed, and three technical replicates of each sample were carried by us. ** means P < 0.01.

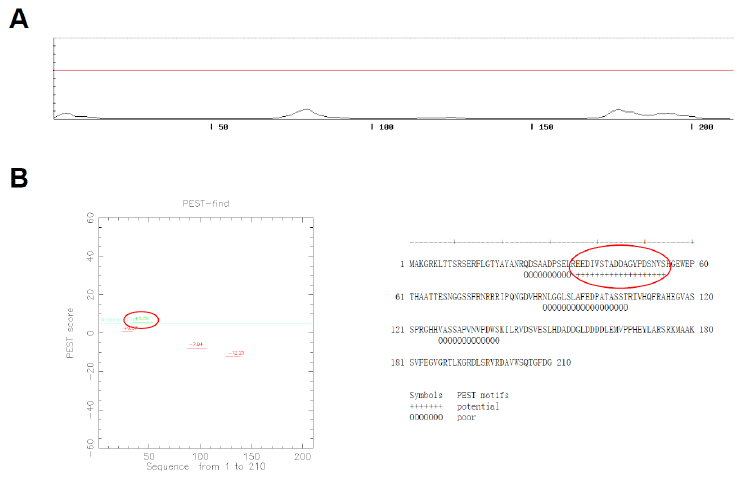

**Supplementary Figure S6. The nuclear localization sequence and the PEST sequence prediction of CiS40-11.**

(A) was the nuclear localization sequence prediction of CiS40-11 with online software NLStradamus (http://www.moseslab.csb.utoronto.ca/). The red line represented prediction cutoff value. (B) was the PEST sequence prediction of CiS40-11 with online software epestfind (https://www.bioinformatics.nl/cgi-bin/emboss/epestfind). The red circles represented the PEST sequence.

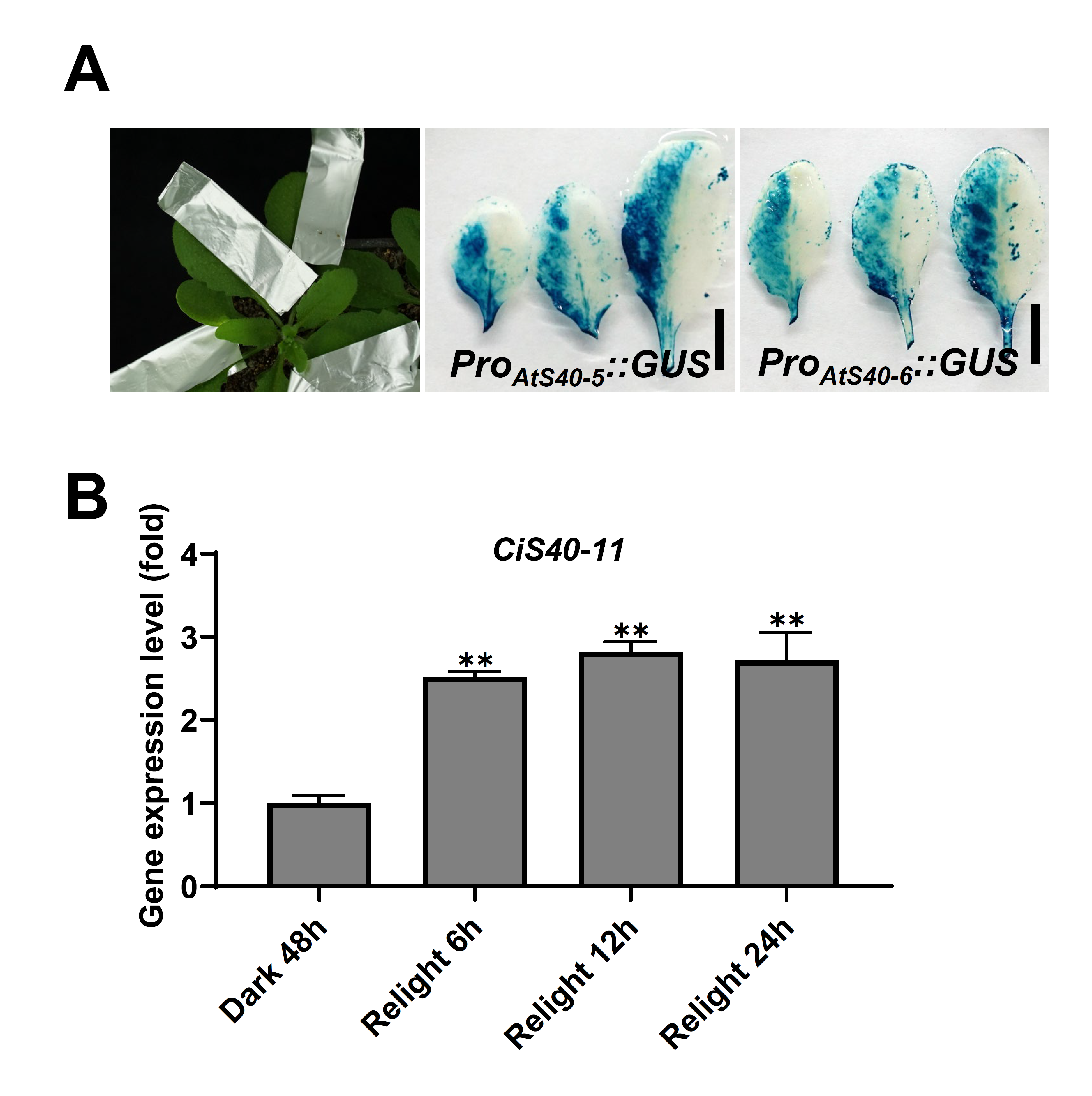

**Supplementary Figure S7. The histochemical staining of *Pro_AtS40-5_::GUS* and *Pro_AtS40-6_::GUS* transgenic *A. thaliana* and the *CiS40-11* expression level detection under light treatment.**

(A) was the histochemical staining of *Pro_AtS40-5_::GUS* and *Pro_AtS40-6_::GUS* transgenic *A. thaliana*. The 2-kb fragment upstream of the *AtS40-5* and *AtS40-6* translation start site was cloned respectively (primer sequences are listed in Supplementary Table S3), and the *Pro_AtS40-5_::GUS* and *Pro_AtS40-6_::GUS* expression vectors were constructed and were transformed into *A. thaliana*. When the transgenic plants had grown about 4 weeks, wrapped the half leaf with tinfoil of the 5^th^, 6^th^, and 7^th^ rosette leaves, as shown in the first panel in **a**. Then plants were cultured under normal condition for three days. Samples were taken and histochemical staining for GUS activity were performed for 12 hours. Pictures were taken after destaining. (B) was the *CiS40-11* expression level detection under light treatment. Four-week-old *C. intermedia* seedlings growing under normal growth condition were subjected to dark treatment for 48 hours, then were exposed to continuous light treatment, and leaves were taken at the indicated time points for qRT-PCR detection. *CiEF1α* was used as the internal reference gene and the expression value was calculated using 2^-ΔΔCT^ method. Seedlings under dark treatment for 48 hours was taken as the control. Three independent biological replicates were performed, and three technical replicates of each sample were carried out. ** means P < 0.01.

**Supplementary Figure S8. The expression level of some *SAGs* changed in the transcriptome and qRT-PCR validation in the OE35 line.**

The gray columns represented *SAGs* expression level changes in the transcriptome, and the black columns represented *SAGs* expression level changes with qRT-PCR detection. Plants had grown for about 5 weeks, *AtEF1α* was used as the internal reference gene and the expression value was calculated using 2^-ΔΔCT^ method. Two independent biological replicates were performed, and three technical replicates of each sample were carried out. The *SAGs* expression level in WT was taken as the control in both the transcriptome and the qRT-PCR analysis.
